# How low can you go? Short-read polishing of Oxford Nanopore bacterial genome assemblies

**DOI:** 10.1101/2024.03.07.584013

**Authors:** George Bouras, Louise M. Judd, Robert A. Edwards, Sarah Vreugde, Timothy P. Stinear, Ryan R. Wick

## Abstract

It is now possible to assemble near-perfect bacterial genomes using Oxford Nanopore Technologies (ONT) long reads, but short-read polishing is still required for perfection. However, the effect of short-read depth on polishing performance is not well understood. Here, we introduce Pypolca (with default and careful parameters) and Polypolish v0.6.0 (with a new careful parameter). We then show that: (1) all polishers other than Pypolca-careful, Polypolish-default and Polypolish-careful commonly introduce false-positive errors at low depth; (2) most of the benefit of short-read polishing occurs by 25× depth; (3) Polypolish-careful never introduces false-positive errors at any depth; and (4) Pypolca-careful is the single most effective polisher. Overall, we recommend the following polishing strategies: Polypolish-careful alone when depth is very low (<5×), Polypolish-careful and Pypolca-careful when depth is low (5–25×), and Polypolish-default and Pypolca-careful when depth is sufficient (>25×).

**Data Summary:** Pypolca is open-source and freely available on Bioconda, PyPI, and GitHub (github.com/gbouras13/pypolca). Polypolish is open-source and freely available on Bioconda and GitHub (github.com/rrwick/Polypolish). All code and data required to reproduce analyses and figures are available at github.com/gbouras13/depth_vs_polishing_analysis. All FASTQ sequencing reads are available at BioProject PRJNA1042815. A detailed list of accessions can be found in Table S1.

## Introduction

Recent advances in the accuracy of Oxford Nanopore Technologies (ONT) sequencing have made it possible to recover near-perfect complete bacterial genomes using only ONT long-read sets. Remaining errors are often fewer than 10 per genome^1^ and usually arise in long homopolymer and DNA modification sites that are difficult to resolve with current ONT data^2,3^. Therefore, using short reads to polish ONT-only assemblies still provides accuracy benefits^4,5^. In this paradigm, the burden shifts from short-read polishers resolving as many errors as possible (i.e. reducing false negatives), to ensuring that polishers do not introduce new errors (i.e. reducing false positives).

For example, consider a hypothetical polisher that fixes 100% of errors in a given genome but also introduces 10 new errors. For an assembly with hundreds of errors (e.g. an ONT-only Trycycler^6^ assembly in 2021), this polisher would greatly improve accuracy. However, for a near-perfect genome with only five errors (e.g. an ONT-only Trycycler assembly in 2024), this polisher would introduce more errors than it fixed.

Given the lack of errors remaining in ONT-only assemblies, another consideration is the short-read depth required for polishing. While 30× coverage is the standard for variant calling in the context of human genomics^7^, the relationship between polishing and sequencing depth in the bacterial context has not been investigated.

In this study, we present three advances. Firstly, we introduce Pypolca, a Python re-implementation of POLCA^8^ with added features. Secondly, we introduce a new version of Polypolish^9^ (v0.6.0). Both Pypolca and Polypolish v0.6.0 include a new ‘--careful’ option that reduces false positives. Finally, we investigate how the performance of Pypolca, Polypolish and other short-read polishing tools vary with short-read depth using a panel of nine deeply sequenced isolate genomes with near-perfect Trycycler assemblies^6^. We show that approximately 25× genome coverage is sufficient to polish almost all ONT-only assembly errors, and that at very low depths (<5×), all short-read polishers other than Pypolca with ‘--careful’ and Polypolish commonly introduce errors.

## Methods

### Pypolca

Pypolca is a Python-only reimplementation of the short-read polisher POLCA^8^. In comparison to POLCA, Pypolca implements a simplified command-line interface, allowing the user to clearly specify input and output files. Additionally, it is available and installable on both macOS and Linux (whereas POLCA can only be run on Linux) and does not require the installation of the MaSuRCA genome assembler like POLCA^10^.

Pypolca, like POLCA, aligns short reads to an assembly using BWA-MEM^11^, processes the alignments with Samtools^12^, runs freebayes^13^ to call variants between the aligned reads and assembly, and then applies well-supported variants to the assembly. POLCA polishes all variants where: (1) at least two aligned reads support the alternate allele, and (2) there are at least twice as many aligned reads supporting the alterative allele compared to the assembly allele. Pypolca retains these thresholds as defaults (‘Pypolca-default’) but allows users to change condition 1 using ‘--min_alt’ and condition 2 using ‘--min_ratio’. The ‘--careful’ flag sets --min_alt to 4 and --min_ratio to 3, which as we show in this study, prevents almost all false positives at low depths without sacrificing error removal. Examples comparing polishing decisions between Pypolca-default and Pypolca-careful are presented in Figure 1.

**Figure 1:**
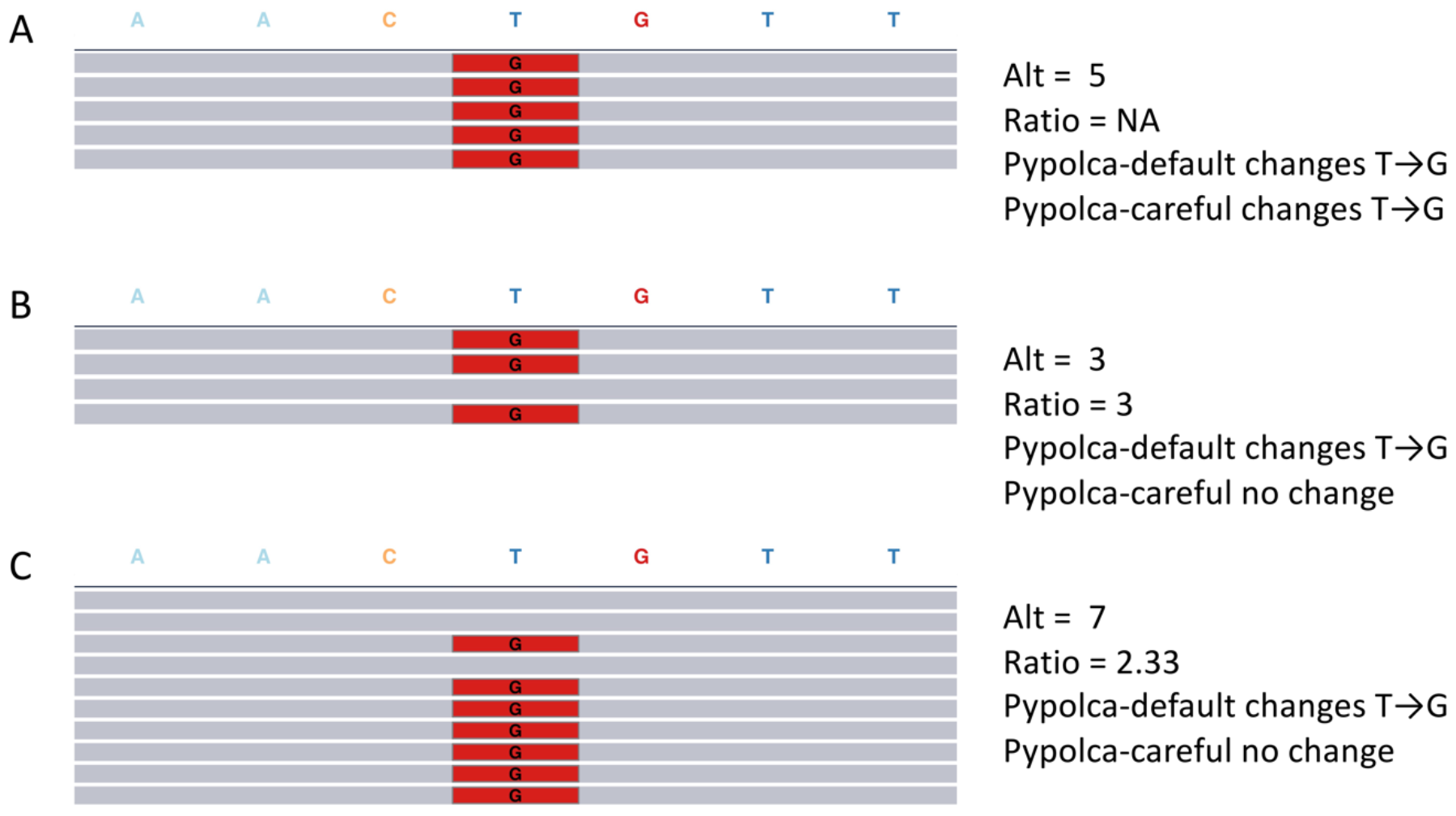
Pypolca polishing decisions at low sequencing depths. Each pileup plot shows simulated reads (represented by horizontal grey bars) aligned to an assembly sequence AAC**T**GTT. Positions that are the same as the assembly are coloured grey, those that differ are coloured red. ‘Alt’ refers to the number of reads aligning to the alternative allele. ‘Ratio’ refers to the number of reads aligning to the alternative allele divided by the number of reads aligning to the assembly allele. **(A)** All 5 aligned reads support the alternative ‘G’ (coloured in red) rather than the assembly ‘T’ at the fifth base in the sequence. Both Pypolca-default and Pypolca-careful will change T→G at this position. **(B)** Three aligned reads have support for ‘G’, while one read supports ‘T’ (grey). In this case Pypolca-default will change T→G, as at least 2 reads support the alternative allele and the alternative-to-assembly ratio is greater than 2. However, because it has only 3 supporting reads (under the threshold of 4), Pypolca-careful will not change this position and leave it as ‘T’. **(C)** Seven aligned reads support ‘G’ while 3 support ‘T’, Pypolca-default will change T→G but Pypolca-careful will not, as the ratio between alternative and reference alleles is 2.33 (under the threshold of 3).

### Polypolish v0.6.0

The Polypolish software and algorithm has been described earlier^9^. As part of this study, we introduce Polypolish v0.6.0, which is now completely implemented in Rust and adds a ‘--careful’ option for low depth polishing. The motivation for --careful comes from polishing repeat regions.

Consider a repeat region in a genome that occurs twice but is not completely identical. At low depths of short-reads, it is possible to obtain reads from only one instance of the repeat. In this scenario Polypolish may ‘polish’ the unsequenced instance of the repeat to be the same as the sequenced instance, thereby introducing a false-positive error.

The ‘--careful’ flag introduced in v0.6.0 forces Polypolish to ignore all reads with multiple alignments. This makes it unable to fix errors in repeats, slightly reducing its ability to fix errors where short-read depth is high, but it ensures that Polypolish will never introduce false-positive polishing errors in repeats regions. However, Polypolish with default parameters is the only alignment-based method that can consistently fix errors in repeat regions and therefore is useful where short-read depth is high (Figure S1).

### Sample Selection

A panel of nine ATCC bacterial genomes were used to compare the effect of depth on short-read polishing of near-perfect ONT-only assembled genomes. The genomes were deeply sequenced with at least 50× short read and long read coverage. They include:

- *Salmonella enterica* ATCC 10708
- *Vibrio cholerae* ATCC 14035
- *Vibrio parahaemolyticus* ATCC 17802
- *Listeria ivanovii* ATCC 19119
- *Escherichia coli* ATCC 25922
- *Campylobacter jejuni* ATCC 33560
- *Campylobacter lari* ATCC 35221
- *Listeria welshimeri* ATCC 35897
- *Listeria monocytogenes* ATCC BAA-679

### Sequencing

DNA extraction was performed with the GenFind V2 kit (Beckman Coulter, Brea, USA). Illumina library preparation was performed with Illumina DNA Prep using quarter reagents (Illumina, San Diego, USA). Short-read whole genome sequencing was performed on an Illumina NextSeq 2000 with a 150bp PE kit (Illumina).

ONT libraries were prepared using the ONT SQK-NBD114-96 and SQK-RBK114-96 kits, and the resultant libraries were sequenced using R10.4.1 MinION flow cells (FLO-MIN114) on a GridION (ONT, Oxford, UK). ONT data was basecalled using the dna_r10.4.1_e8.2_sup@v4.3.0 model with Dorado v0.5.0.

### Benchmarking

ONT-only assemblies were created using Trycycler v0.5.4 and reoriented to begin with a consistent starting position using Dnaapler^14^ v0.7.0 as described by us in previous studies using this data^1,15^. The number of errors in the ONT-only assemblies (compared to our previously published method for generating robust and curated bacterial reference genome assemblies^16^) ranged from 0 to 18 total errors, for a total of 37 errors across all 9 genomes. Medaka polishing was not performed, as we found it to have a net negative impact on accuracy (Table S2).

Short-read FASTQs were first quality controlled with fastp^17^ v0.23.4. Short-read FASTQs from genome were subsampled using the ‘seqtk sample’ command from seqtk^18^ at 0.1× depth intervals from 0.1× estimated genome coverage up to 50×, yielding 500 subsampled read sets.

For every subsampled read set, the Trycycler reference genome was polished using the following polishing tools (using default parameters except where specified):

1. Polypolish v0.6.0 (denoted as ‘Polypolish-default’)
2. Polypolish v0.6.0 with ‘--careful’ (denoted as ‘Polypolish-careful’)
3. Pypolca v0.3.0 (denoted as ‘Pypolca-default’)
4. Pypolca v0.3.0 with ‘--careful’ (denoted as ‘Pypolca-careful’) 5. HyPo^19^ v1.0.3
5. FMLRC2^20^ v0.1.8
6. NextPolish^21^ v1.4.1
7. Pilon^22^ v1.24

## Results

We tested all polishers at every interval of short-read coverage from 0.1× to 50× estimated genome coverage. We first compared remaining error counts to the total Trycycler ONT-only error count of 37 across the nine genomes. Every polisher other than Polypolish-careful had a total error count greater than 37 in at least one interval tested (Figure 2; Figure S2).

**Figure 2:**
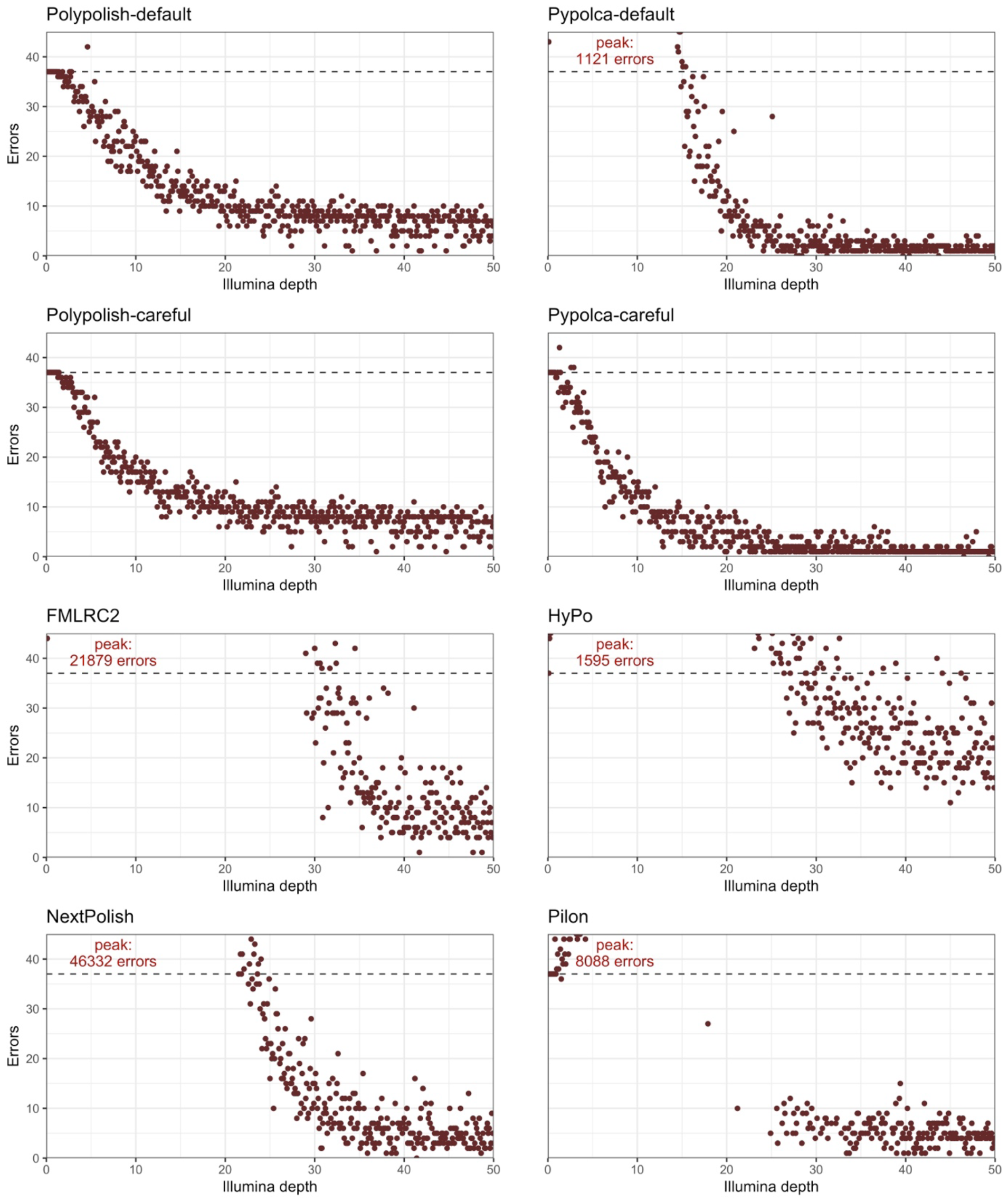
total errors by depth per polisher. Each plot shows the total number of errors remaining in the nine reference genomes at each interval from 0.1× to 50× depth (x-axis) for the eight polishers tested. The dashed blue lines represent the total Trycycler ONT-only assembly error count of 37 errors. Points above this indicate that the polisher has decreased total accuracy, below that the polisher has increased total accuracy. The y-axes for the plots are limited at 45 total errors, with the peak error count labelled in the top left if it exceeds 45. See Figure S2 for the plots with unrestricted y-axes.

Polypolish-default and Pypolca-careful only rarely exceeded 37 total errors (1/500 and 3/500 intervals) and in these cases, the total error counts were only slightly higher (42 total errors for Polypolish-default at 4.6× coverage, and 42, 38 and 38 for Pypolca-careful at 1.3×, 2.6× and 2.9× coverage respectively).

All other polishers routinely introduced many errors at low depths (Figure 2; Figure S2). The maximum number of total errors of these polishers ranged from 1121 (Pypolca-default at 3.3× coverage) to 46332 (NextPolish at 1.6× coverage).

The depths at which these polishers did not decrease accuracy varied. The lowest depth (above 2×) that Pypolca-default had 37 or fewer errors was 14.9×, 17.9× for Pilon, 21.5× for NextPolish, 26.4× for HyPo, 29.1× for FMLRC2. The highest coverage where the polishers had 37 or more errors ranged from 15.4× for Pypolca-default, 24.0× for NextPolish, 34.5× for FMLRC2, 43.5× for HyPo and 45.9× for Pilon.

Next, we analysed the introduction of errors on a per-genome basis out of the nine tested. Polypolish-careful was the most conservative polisher in this analysis. It never decreased accuracy (i.e. increased the total error count compared to the Trycycler ONT-only assembly) in any genome at any interval (Table 1; Figure S3). It was followed by Pypolca-careful, which decreased accuracy in at least one genome in 45/500 intervals, in 51/4500 genomes overall and never above 16.1× depth, and Polypolish-default (103/500 intervals, 147/4500 genomes and never above 21.0× depth). All other polishers commonly decreased accuracy in at least one out of the nine genomes in an interval, even at higher depths (Table 1; Figure S3).

**Table 1:** total number of genomes and intervals where polishing decreased accuracy (i.e. total error count was increased compared to the Trycycler assembly).

| Polisher | Intervals with decreased accuracy in at least 1 genome (n= 500) | Total genomes with decreased accuracy (n= 4500) |
| --- | --- | --- |
| **FMLRC2** | 430 | 2671 |
| **HyPo** | 499 | 3097 |
| **NextPolish** | 386 | 2189 |
| **Pilon** | 324 | 1331 |
| **Polypolish-default** | 103 | 147 |
| **Polypolish-careful** | 0 | 0 |
| **Pypolca-default** | 221 | 1374 |
| **Pypolca-careful** | 45 | 51 |

We then focused on performance of polishers above 25× depth (Table S3). Three polishers (Pypolca-default 7 times, Pypolca-careful 2 times and NextPolish 1 time) had at least one interval where all errors were resolved. Every other polisher had at least one interval where only 1 error was remaining, other than HyPo (minimum of 11 remaining errors).

Pypolca-careful had the lowest mean and median error count, followed by Pypolca-default (Table S3). Pypolca-careful had a median error of 1, which was a difficult to polish error in a long homopolymer on a *S. enterica* plasmid (Figure S4). Polypolish-default, Polypolish-careful and NextPolish never made ONT-only assemblies worse above 25×, though Polypolish in both modes had significantly lower variance than NextPolish. HyPo and FMLRC2 on average improved assemblies, though not always, while Pilon on average made assemblies worse.

Due to the strong and consistent performance of Pypolca and Polypolish, we then compared pairwise combinations of Pypolca-default, Pypolca-careful, Polypolish-default and Polypolish-careful run sequentially. Overall, we found the results to be concordant with the single-tool polishing results, with combinations including Pypolca-careful performing the best overall. The order of polishers had no impact of the results (Table S4; Figure S5).

A combination of Pypolca-careful and Polypolish depending on estimated short-read depth has been implemented in our automated assembly tool Hybracter from v0.7.0 ^15^. Consistent with these results, when the automated ONT-only assembly did not have structural errors ^23^, running Hybracter on the benchmarked genomes was consistently able to produce polished assemblies that were error-free above 25× depth, and sometimes even at lower depth (Figure S6). Short-read polishing almost always improved Hybracter assemblies. However, improvements in assembly quality were frequently not reflected in higher reference-free ALE scores, suggesting that where near-perfect ONT-only assemblies are polished as in this study, using ALE has limited utility (Figure S7).

Finally, we binned all intervals for every 5× depth increase for the three best performing polishers (Pypolca-careful, Polypolish-default and Polypolish-careful). We calculated the mean number of total remaining errors in each bin. Short-read polishing improved assemblies for every short-read depth bin (Table 2). Most of the benefit of polishing is seen at depths below 25×, after which the benefits of extra depth on polishing performance are minor.

**Table 2:** mean remaining errors at all depth intervals within every 5× depth bin from 0.1× to 50× for the three best polishers (Polypolish-careful, Polypolish-default, Pypolca-careful)

| Polisher | 0.1–<br>5× | 5.1–<br>10× | 10.1–<br>15× | 15.1–<br>20× | 20.1–<br>25× | 25.1–<br>30× | 30.1–<br>35× | 35.1–<br>40× | 40.1–<br>45× | 45.1–<br>50× |
| --- | --- | --- | --- | --- | --- | --- | --- | --- | --- | --- |
| **Polypolish-careful** | 33.48 | 20.10 | 13.70 | 11.52 | 8.98 | 8.56 | 7.66 | 7.28 | 6.50 | 6.44 |
| **Polypolish-default** | 34.26 | 24.12 | 15.68 | 11.98 | 8.86 | 8.46 | 7.50 | 7.14 | 6.28 | 6.32 |
| **Pypolca-careful** | 31.90 | 15.76 | 8.50 | 5.02 | 3.60 | 1.96 | 1.94 | 1.80 | 1.16 | 1.24 |

## Discussion

In this study, we show that at low depths (<25×), many widely used short-read polishing tools frequently introduce, rather than reduce errors in bacterial genome assemblies. The only exceptions are Polypolish-careful, which never introduced an error, Polypolish-default and Pypolca-careful, which rarely introduced errors (51/4500 samples and 147/4500 samples respectively). We also show in this study that most of the utility in short-read polishing occurs by approximately 25× average read depth across the genome.

While it would be unusual to have less than 25× average short-read depth across the genome for single isolate genome assemblies, there are scenarios where this concern applies even where overall sequencing depth is high, necessitating the use of conservative polishers. The first is for genomes, or regions within genomes, that have extreme GC content. It has been shown that extremes in GC-content can lead to issues in PCR-amplification steps that lead to low read depth in GC-rich regions with short-read sequencing ^24,25^. In such regions, short-read polishers that are not conservative may introduce errors even where the overall genome coverage is high. Regions of local low read depth may also arise in genomes with neutral GC-content, particularly where transposase-based library preparation kits are used^26^.

Another scenario is polishing ONT metagenomic assemblies^27^. Assembling metagenome assembled genomes (MAGs) is a difficult problem due to variations in sequencing depth and community composition^28,29^. While ONT-only metagenomic assemblies are improving ^27^, short-read polishing is still commonly recommended to improve genome completeness and protein-coding annotation of MAGs and contigs^30,31^. As even deep sequencing can rarely, if ever, recover the full extent of diversity^32^, many metagenomic assembled contigs will not constitute complete genomes and have low coverage (under 25×). If these contigs are polished with short-reads with similarly low coverage, using short-read polishers may introduce errors in these contigs. Further, long-read and short-read methods may not recover the same populations from within a metagenome^31^, reducing coverage even further and emphasising the need to use a conservative short-read polisher.

Our analysis showed that Pypolca-careful was the single best polisher tested, with no improvements when combined with the next best short-read polisher (Polypolish-careful) in our benchmarked dataset. It is also known that Medaka polishing of ONT assemblies can introduce errors to near-perfect genomes and is not recommended for Q20+ Nanopore assemblies data^1,33^, which also occurs in this study (see Table S2). Therefore, for the bacterial isolate use-case with Q20+ Nanopore data, we recommend no long-read polishing then short-read polishing with Pypolca-careful in all scenarios other than where depth is extremely low (<5× coverage), or if avoiding false-positive changes is vital. If a false-positive polishing rate of 0 is required, we recommend short-read polishing with Polypolish-careful only, as it never introduced a false-positive change in the 4500 samples tested in this study.

At low depths (5–25×), we recommend combining Pypolca-careful with Polypolish-default. Though our benchmarking did not show any improvements of this combination compared to just running Pypolca-careful (Table S4), we showed that Polypolish-careful never makes an assembly worse. In other datasets, this combination could show some improvements compared to Pypolca-careful alone.

At higher depths (above 25×), we recommend Polypolish-default combined with Pypolca-careful. While this was no better than running Pypolca-careful in this dataset (Table S4), adding Polypolish-default could enable the polishing of errors in repeat regions that Pypolca-careful could miss (Figure S1). We have incorporated the above recommendations into Hybracter’s^15^ polishing logic from v0.7.0 and show that is it possible to consistently recover automated perfect genome assemblies when short-read depth is 25× or higher (Figure S6).

If high-depth short-reads are available and manual curation by inspecting manual read alignments with IGV^34^ is possible, some alternatives may be to run Pypolca with relaxed settings (for example --min_ratio 1.5), to fix errors where short reads may be inconsistent. Additionally, FMLRC2’s alignment free approach may be a good option to polish errors that are difficult to fix with alignment-based approaches (Figure S1), but manual curation should be used to screen for false positive changes.

## Conclusion

In this study, we introduce Pypolca (both -default and -careful) and Polypolish-careful as short-read polishing tools. By testing a panel of nine bacterial genomes assembled using Q20+ Nanopore followed with short-read polishing, we show that short-read depth has a significant impact on polishing performance of bacterial isolate genomes. All polishers other than Pypolca-careful, Polypolish-default and Polypolish-careful introduced many false-positive errors at low depth. We show that Pypolca-careful is the best polisher overall and that Polypolish-careful never introduces false-positive errors. We recommend targeting a short-read sequencing depth of 25× or greater when creating bacterial genome assemblies.

Further studies are required to assess the effect of short-read depth on polishing in other contexts, such as extreme GC-content, metagenomes, or eukaryotic genomes.

## Acknowledgements

This research was performed in part at Doherty Applied Microbial Genomics, Department of Microbiology and Immunology, University of Melbourne at the Peter Doherty Institute for Infection and Immunity

## References

1. Wick, R. ONT-only accuracy: 5 kHz and Dorado. Ryan Wick’s bioinformatics blog https://rrwick.github.io/2023/10/24/ont-only-accuracy-update.html 10.5281/zenodo.10038672 (2023).

2. Wick, R. R. & Holt, K. E. Benchmarking of long-read assemblers for prokaryote whole genome sequencing. Preprint at 10.12688/f1000research.21782.4 (2021).

3. Delahaye, C. & Nicolas, J. Sequencing DNA with nanopores: Troubles and biases. PLOS ONE 16, e0257521 (2021).

4. Lerminiaux, N., Fakharuddin, K., Mulvey, M. R. & Mataseje, L. Do we still need Illumina sequencing data?: Evaluating Oxford Nanopore Technologies R10.4.1 flow cells and v14 library prep kits for Gram negative bacteria whole genome assemblies. 2023.09.25.559359 Preprint at 10.1101/2023.09.25.559359 (2023).

5. Sanderson, N. D. et al. Evaluation of the accuracy of bacterial genome reconstruction with Oxford Nanopore R10.4.1 long-read-only sequencing. 2024.01.12.575342 Preprint at 10.1101/2024.01.12.575342 (2024).

6. Wick, R. R. et al. Trycycler: consensus long-read assemblies for bacterial genomes. Genome Biology 22, 266 (2021).

7. Bentley, D. R. et al. Accurate whole human genome sequencing using reversible terminator chemistry. Nature 456, 53–59 (2008).

8. Zimin, A. V. & Salzberg, S. L. The genome polishing tool POLCA makes fast and accurate corrections in genome assemblies. PLOS Computational Biology 16, e1007981 (2020).

9. Wick, R. R. & Holt, K. E. Polypolish: Short-read polishing of long-read bacterial genome assemblies. PLOS Computational Biology 18, e1009802 (2022).

10. Zimin, A. V. et al. The MaSuRCA genome assembler. Bioinformatics 29, 2669–2677 (2013).

11. Li, H. Aligning sequence reads, clone sequences and assembly contigs with BWA-MEM. Preprint at 10.48550/arXiv.1303.3997 (2013).

12. Li, H. et al. The Sequence Alignment/Map format and SAMtools. Bioinformatics 25, 2078–2079 (2009).

13. Garrison, E. & Marth, G. Haplotype-based variant detection from short-read sequencing. Preprint at 10.48550/arXiv.1207.3907 (2012).

14. Bouras, G., Grigson, S. R., Papudeshi, B., Mallawaarachchi, V. & Roach, M. J. Dnaapler: A tool to reorient circular microbial genomes. Journal of Open Source Software 9, 5968 (2024).

15. Bouras, G. et al. Hybracter: Enabling Scalable, Automated, Complete and Accurate Bacterial Genome Assemblies. 2023.12.12.571215 Preprint at 10.1101/2023.12.12.571215 (2023).

16. Wick, R. R., Judd, L. M. & Holt, K. E. Assembling the perfect bacterial genome using Oxford Nanopore and Illumina sequencing. PLOS Computational Biology 19, e1010905 (2023).

17. Chen, S., Zhou, Y., Chen, Y. & Gu, J. fastp: an ultra-fast all-in-one FASTQ preprocessor. Bioinformatics 34, i884–i890 (2018).

18. Li, H. seqtk: a fast and lightweight tool for processing sequences in the FASTA or FASTQ format. https://github.com/lh3/seqtk.

19. Kundu, R., Casey, J. & Sung, W.-K. HyPo: Super Fast & Accurate Polisher for Long Read Genome Assemblies. 2019.12.19.882506 Preprint at 10.1101/2019.12.19.882506 (2019).

20. Mak, Q. X. C., Wick, R. R., Holt, J. M. & Wang, J. R. Polishing De Novo Nanopore Assemblies of Bacteria and Eukaryotes With FMLRC2. Molecular Biology and Evolution 40, msad048 (2023).

21. Hu, J., Fan, J., Sun, Z. & Liu, S. NextPolish: a fast and efficient genome polishing tool for long-read assembly. Bioinformatics 36, 2253–2255 (2020).

22. Walker, B. J. et al. Pilon: An Integrated Tool for Comprehensive Microbial Variant Detection and Genome Assembly Improvement. PLOS ONE 9, e112963 (2014).

23. Wick, R. R. & Bouras, G. A tale of two misassemblies. Ryan Wick’s bioinformatics blog https://rrwick.github.io/2024/02/15/misassemblies.html 10.5281/zenodo.10662704 (2024).

24. Benjamini, Y. & Speed, T. P. Summarizing and correcting the GC content bias in high-throughput sequencing. Nucleic Acids Research 40, e72 (2012).

25. Chen, Y.-C., Liu, T., Yu, C.-H.Chiang, T.-Y. & Hwang, C.-C. Effects of GC Bias in Next-Generation-Sequencing Data on De Novo Genome Assembly. PLOS ONE 8, e62856 (2013).

26. Segerman, B., Ástvaldsson, Á., Mustafa, L., Skarin, J. & Skarin, H. The efficiency of Nextera XT tagmentation depends on G and C bases in the binding motif leading to uneven coverage in bacterial species with low and neutral GC-content. Frontiers in Microbiology 13, (2022).

27. Sereika, M. et al. Oxford Nanopore R10.4 long-read sequencing enables the generation of near-finished bacterial genomes from pure cultures and metagenomes without short-read or reference polishing. Nat Methods 19, 823–826 (2022).

28. Cattonaro, F., Spadotto, A., Radovic, S. & Marroni, F. Do you cov me? Effect of coverage reduction on metagenome shotgun sequencing studies. F1000Res 7, 1767 (2020).

29. Benoit, G. et al. High-quality metagenome assembly from long accurate reads with metaMDBG. Nat Biotechnol 1–6 (2024) doi:10.1038/s41587-023-01983-6.

30. Liu, L., Yang, Y., Deng, Y. & Zhang, T. Nanopore long-read-only metagenomics enables complete and high-quality genome reconstruction from mock and complex metagenomes. Microbiome 10, 209 (2022).

31. Cook, R. et al. Nanopore and Illumina Sequencing Reveal Different Viral Populations from Human Gut Samples. 2023.11.24.568560 Preprint at 10.1101/2023.11.24.568560 (2023).

32. van der Walt, A. J. et al. Assembling metagenomes, one community at a time. BMC Genomics 18, 521 (2017).

33. Wick, R. ONT-only accuracy with R10.4.1. Ryan Wick’s bioinformatics blog https://rrwick.github.io/2023/05/05/ont-only-accuracy-with-r10.4.1.html 10.5281/zenodo.7898220 (2023).

34. Robinson, J. T. et al. Integrative Genomics Viewer. Nat Biotechnol 29, 24–26 (2011).

