## Supplementary Figures for "How low can you go? Short-read polishing of Oxford Nanopore bacterial genome assemblies"

### Main Figures

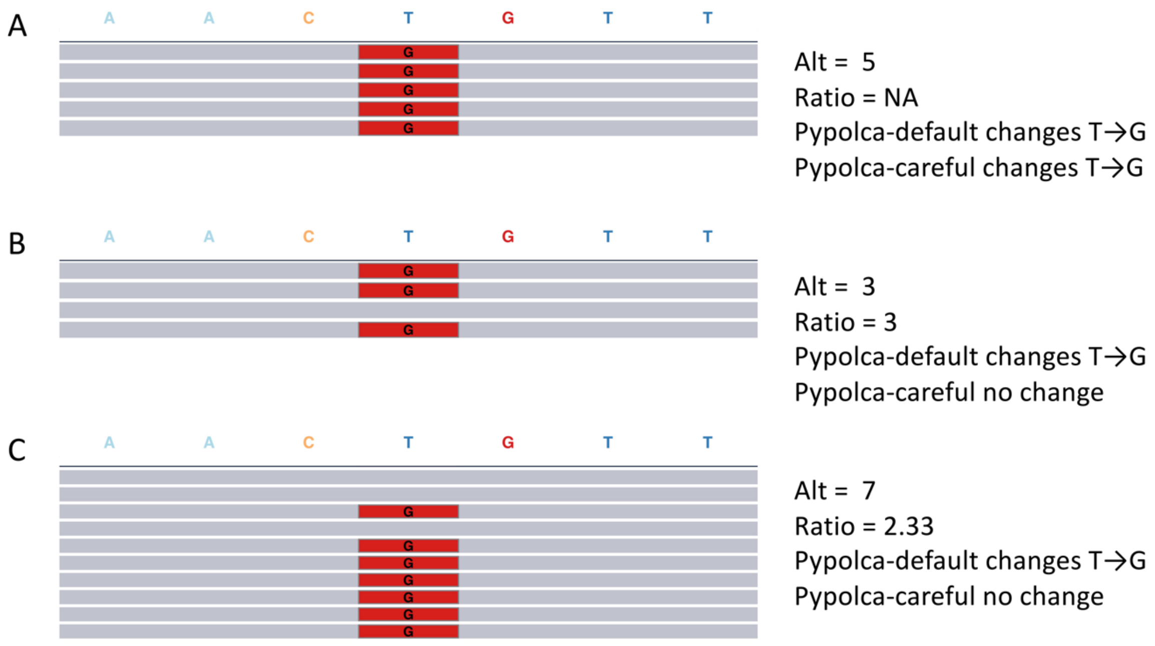


**Figure 1: Pypolca polishing decisions at low sequencing depths.** Each pileup plot shows simulated reads (represented by horizontal grey bars) aligned to an assembly sequence AAC**T**GTT. Positions that are the same as the assembly are coloured grey, those that differ are coloured red. 'Alt' refers to the number of reads aligning to the alternative allele. 'Ratio' refers to the number of reads aligning to the alternative allele divided by the number of reads aligning to the assembly allele. (A) All 5 aligned reads support the alternative 'G' (coloured in red) rather than the assembly 'T' at the fifth base in the sequence. Both Pypolca-default and Pypolca-careful will change T→G at this position. (B) Three aligned reads have support for 'G', while one read supports 'T' (grey). In this case Pypolca-default will change T→G, as at least 2 reads support the alternative allele and the alternative-to-assembly ratio is greater than 2. However, because it has only 3 supporting reads (under the threshold of 4), Pypolca-careful will not change this position and leave it as 'T'. (C) Seven aligned reads support 'G' while 3 support 'T', Pypolca-default will change T→G but Pypolca-careful will not, as the ratio between alternative and reference alleles is 2.33 (under the threshold of 3).

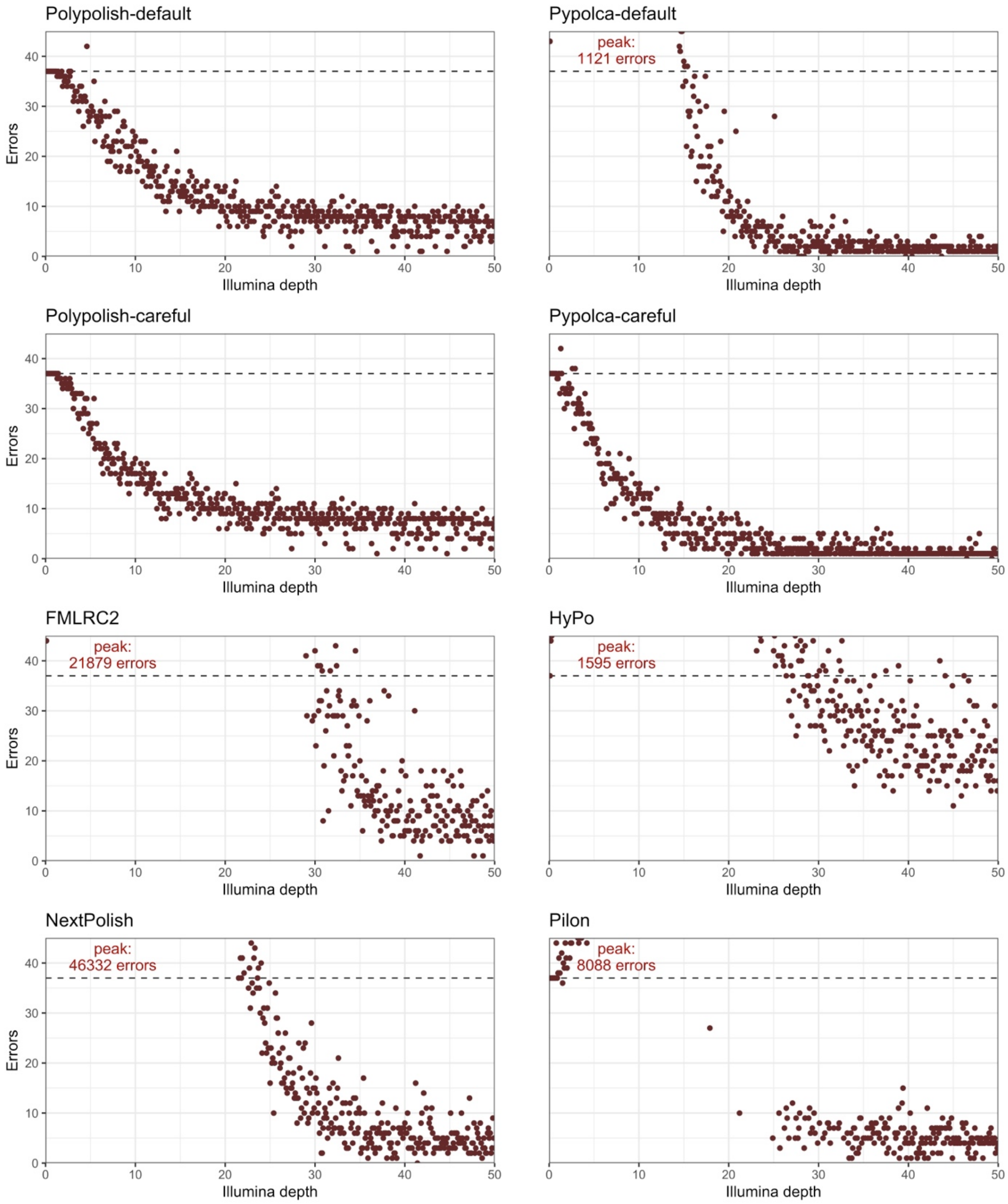


**Figure 2: total errors by depth per polisher**. Each plot shows the total number of errors remaining in the nine reference genomes at each interval from 0.1× to 50× depth (x-axis) for the eight polishers tested. The dashed blue lines represent the total Trycycler long-read only assembly error count of 37 errors. Points above this indicate that the polisher has decreased total accuracy, below that the polisher has increased total accuracy. The y-axes for the plots are limited at 45 total errors, with the peak error count labelled in the top left if it exceeds 45. See Figure S2 for the plots with unrestricted y-axes.

### Supplementary Figures

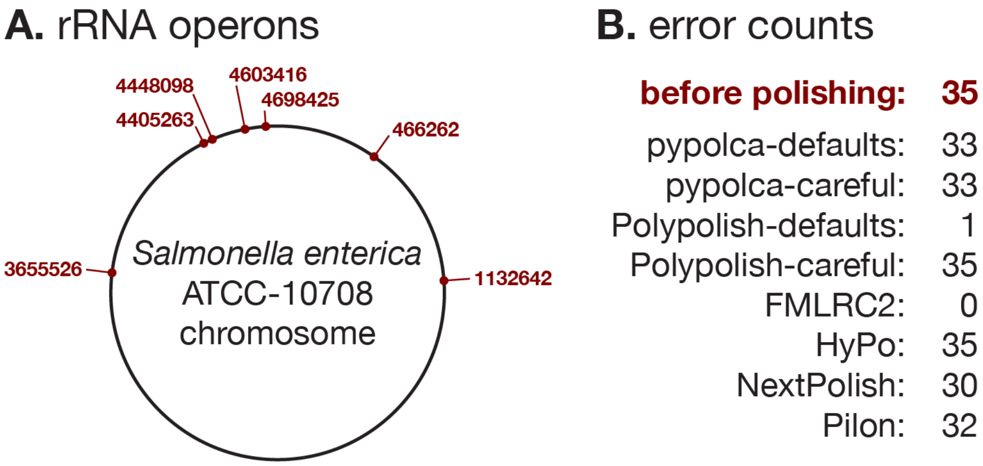


**Figure S1: Polypolish-default can fix errors in repeats other alignment-based tools cannot.** There was only one error out of 37 from the benchmarked Trycycler assemblies near a repeat region (*C. lari* at position 491989) and that error was right at the edge of a repeat. Therefore, to illustrate the continuing utility of Polypolish-default in repeat regions, we added five random substitution errors to each of the seven copies of the ~5.5kbp rRNA operon in *S. enterica* ATCC 10708. This created a total of 35 simulated errors per genome (A). Each polishing tool was then run on the error-containing genome. Polypolish-default was the only alignment-based tool that could correct most errors, with only 1/35 remaining (B). All other alignment-based polishers did not correct many errors (30-35 remaining errors), although FMLRC2, a non-alignment-based method, could correct every error.

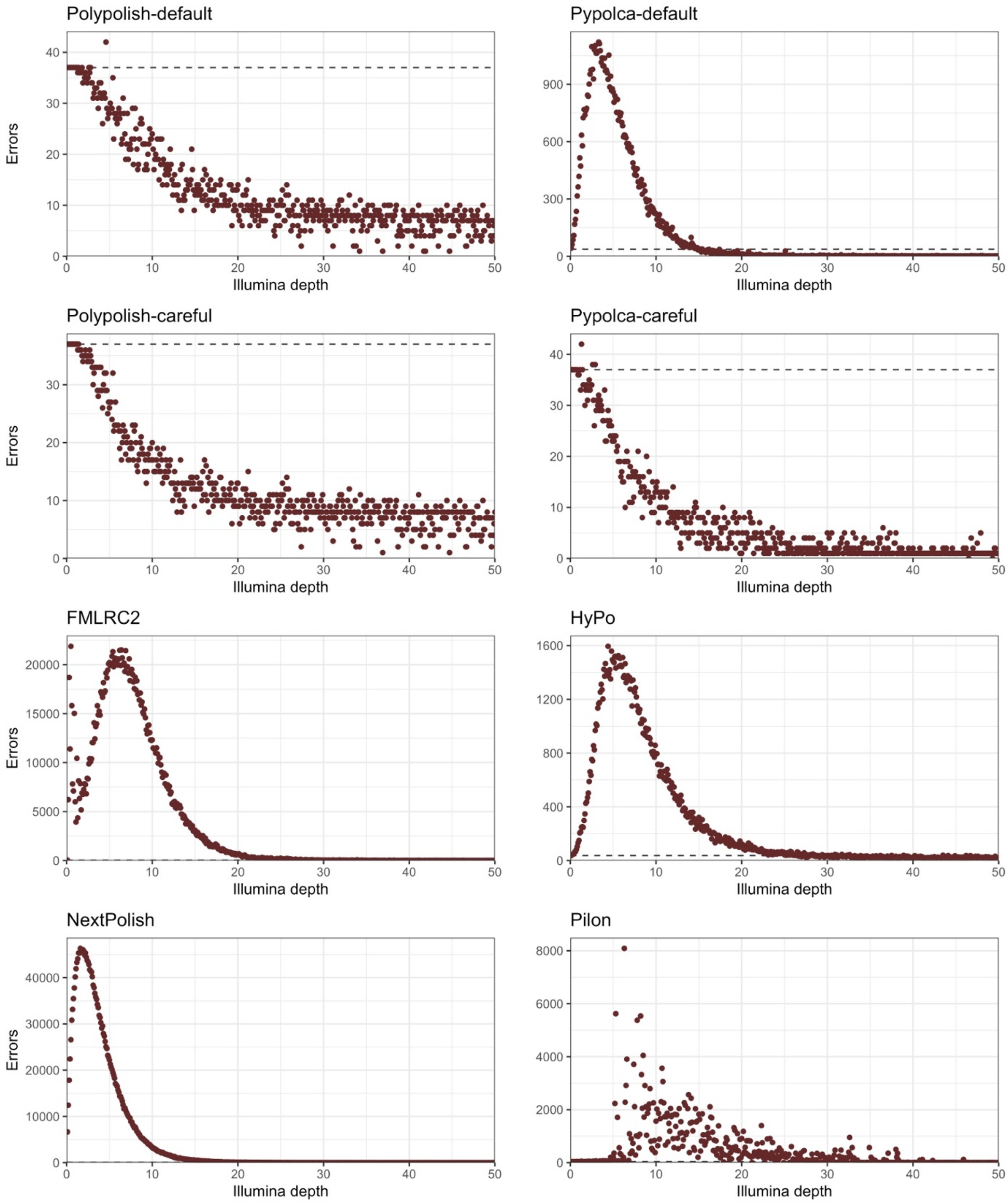


**Figure S2: total errors by depth per polisher with unrestricted y-axes**. This figure is the same as Figure 2 but with full range on the y-axes. Each plot shows the total number errors remaining in the 9 reference genomes at each interval from 0.1× to 50× depth (x-axis) for the 8 single polishers tested. The dashed blue lines represent the total Trycycler long-read only assembly error count of 37 errors. Points above this indicate that the polisher has decreased accuracy, below that the polisher has increased accuracy. The y-axes for the plots are not limited.

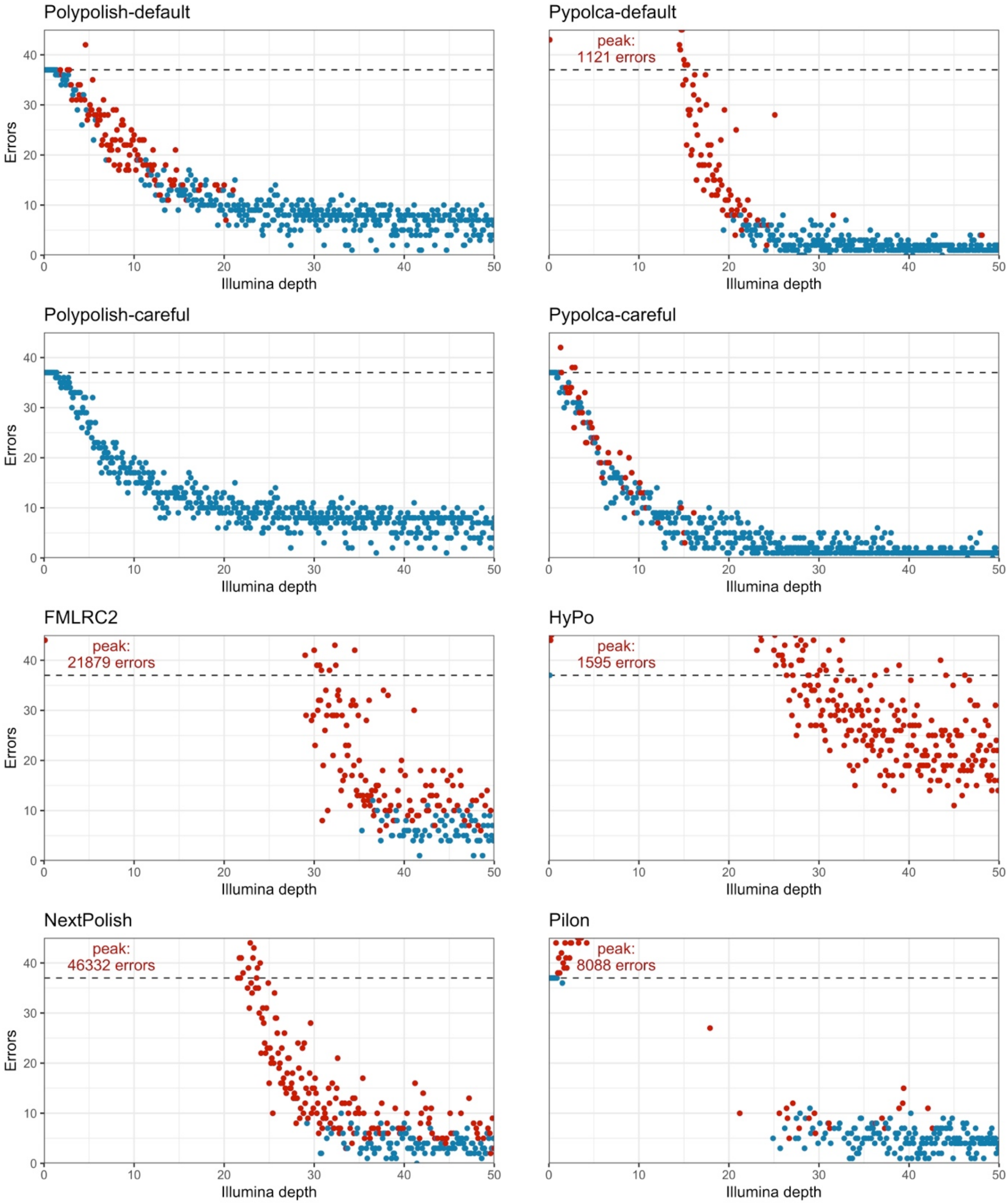


**Figure S3: total errors by depth per polisher coloured where accuracy is decreased in at least one genome**. This figure is the same as Figure 2 but with additional information shown in the colour of the points. Red points indicate all intervals where there were more total errors compared to the Trycycler baseline in at least one out of the nine genomes tested. Blue points indicate intervals where there all nine polished genomes had equal or fewer errors than the Trycycler assemblies.

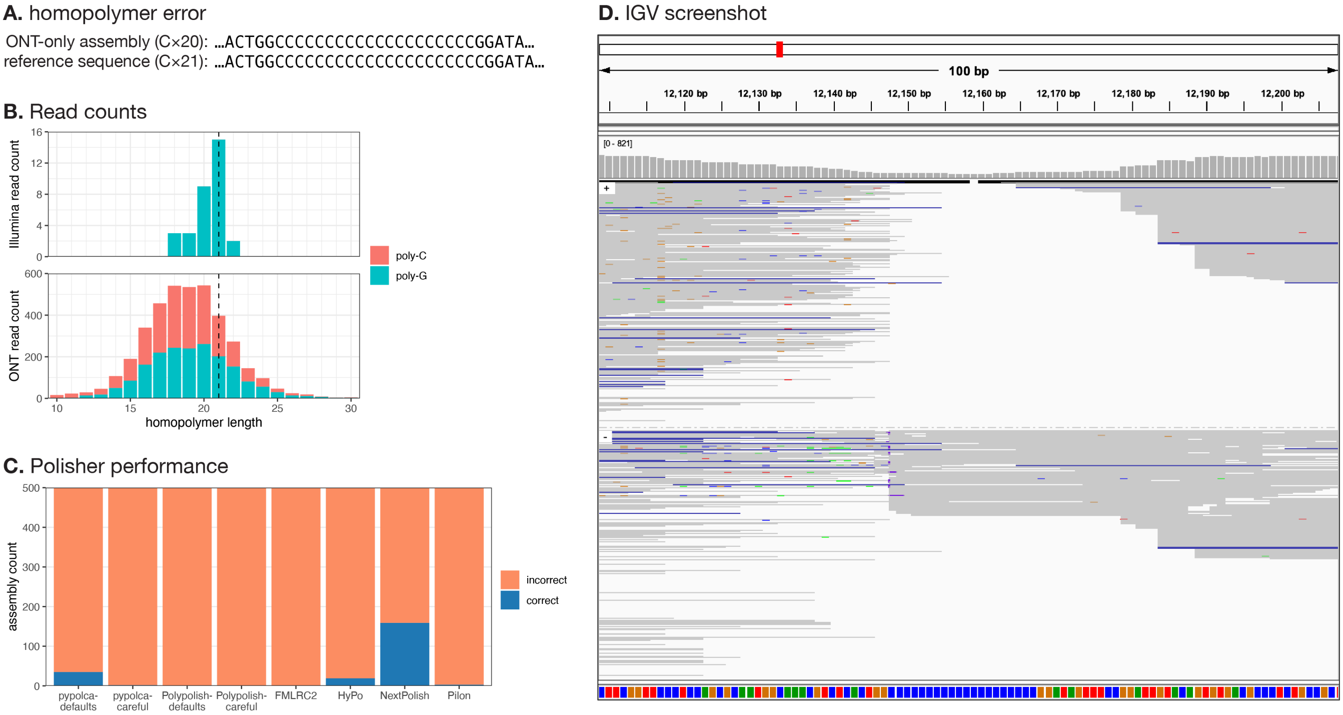


**Figure S4**: **long homopolymer in *Salmonella enterica* ATCC-10708.** (A) This genome's plasmid contains a long homopolymer at position 12148, and there is a discrepancy between the ONT-only assembly (C×20) and the polished reference sequence (C×21). (B) Illumina and ONT reads counts with exact matches to the homopolymer plus five bp of adjacent sequence on both sides. The dashed line at (C×21). indicates the homopolymer length in the reference sequence. For Illumina reads, there were 32 matching reads, (C×21). was the most common homopolymer length, and all matching reads were on the G-strand. For ONT reads, there were 3895 matching reads, (C×18) to (C×20) were the most common homopolymer lengths, and matching reads occurred on both strands. These results create uncertainty in the true homopolymer length for this genome, and the (C×21) in the reference sequence is only a best guess. (C) Assuming that (C×21) is the correct homopolymer length, this plot shows assembly counts with the correct (blue) and incorrect (orange) length at this homopolymer, for each polisher tested. NextPolish performed best, but none of the polishers were able to reliably fix this error. (D) Integrative Genomics Viewer (IGV) screenshot of the relevant region of the plasmid, with Illumina read alignments grouped by strand. This was generated using the full set of Illumina reads (367× depth).

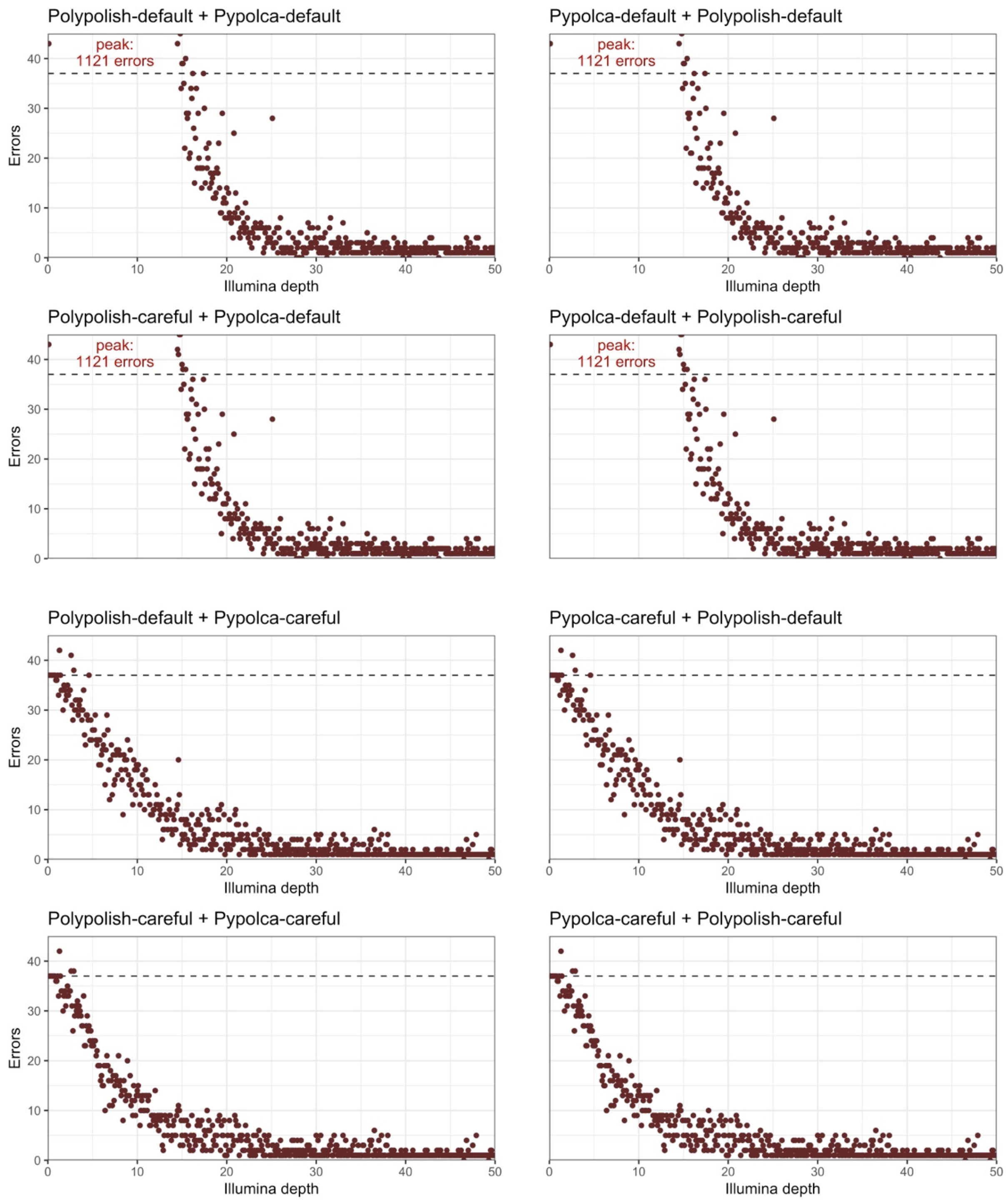


**Figure S5: total errors in Polypolish + Pypolca combinations**. Each plot shows the total number errors remaining in the nine reference genomes at each interval from 0.1× to 50× depth (x-axis) for the eight sequential combinations of Pypolca-default, Pypolca-careful, Polypolish-default and Polypolish-careful tested. The first polisher is presented first in each panel's title, followed by the second polisher. The dashed blue lines represent the total Trycycler long-read only assembly error count of 37 errors. Points above this indicate that the polisher has decreased accuracy and below that the polisher has increased accuracy. The y-axes for the plots are limited at 45 errors, with the peak error count labelled in the top left if it exceeds 45.

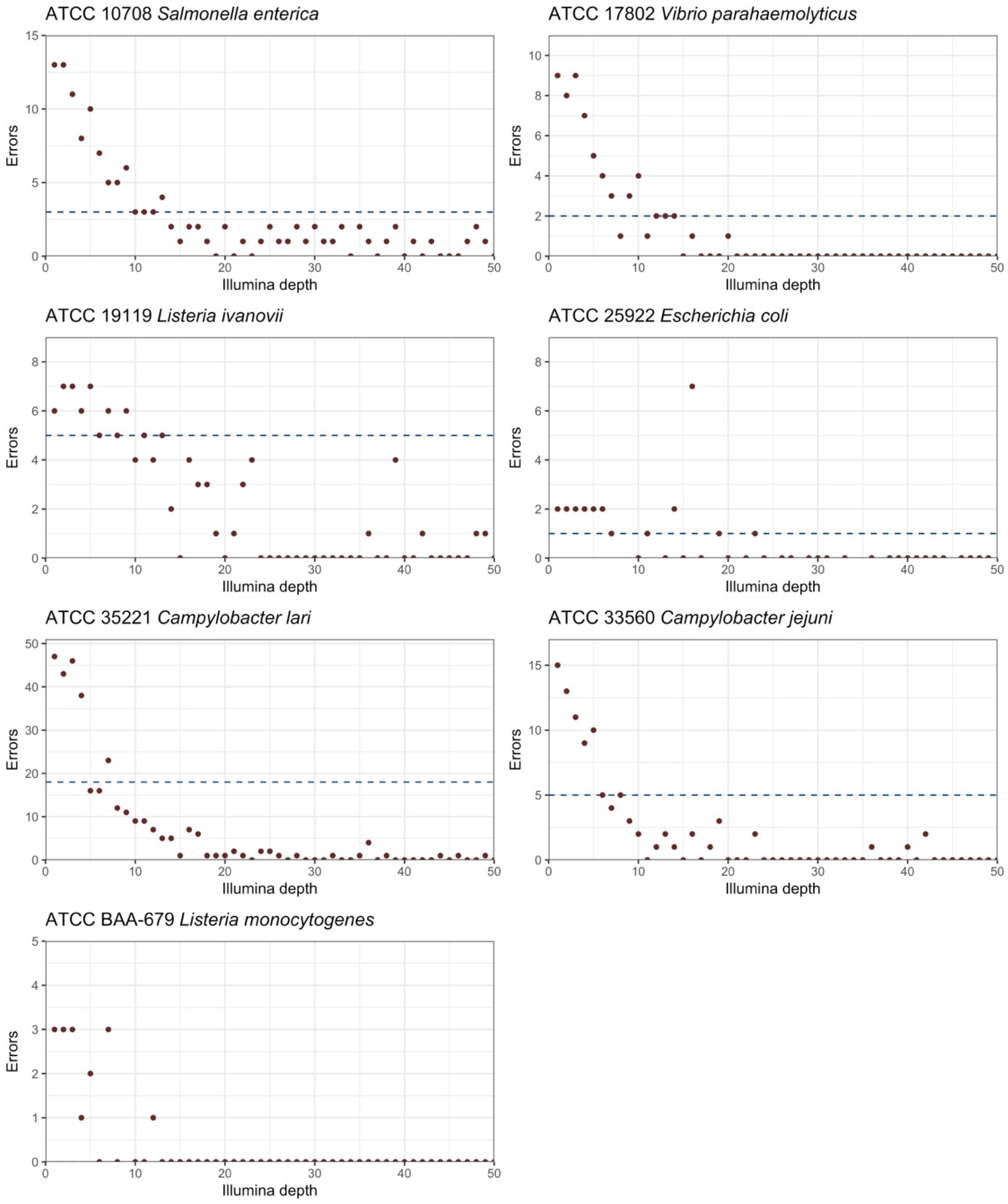


**Figure S6: Hybracter v0.7.0 with short-read polishing errors per genome.** For each of the seven reference genomes without errors and significant non-determinism in their automated long-read assemblies, this Figure shows the number of errors remaining after assembly with Hybracter v0.7.0. Each genome was tested at each interval from 1× to 50× depth (x-axis). The dashed blue lines represent the Trycycler long-read only assembly error count for each genome, with ATCC BAA-679 *L. monocytogenes* having 0 errors in its Trycycler long-read only assembly. Points above this indicate that Hybracter with short-read polishing performs worse than the long-read only Trycycler assemblies, while below indicated that Hybracter has performed better.

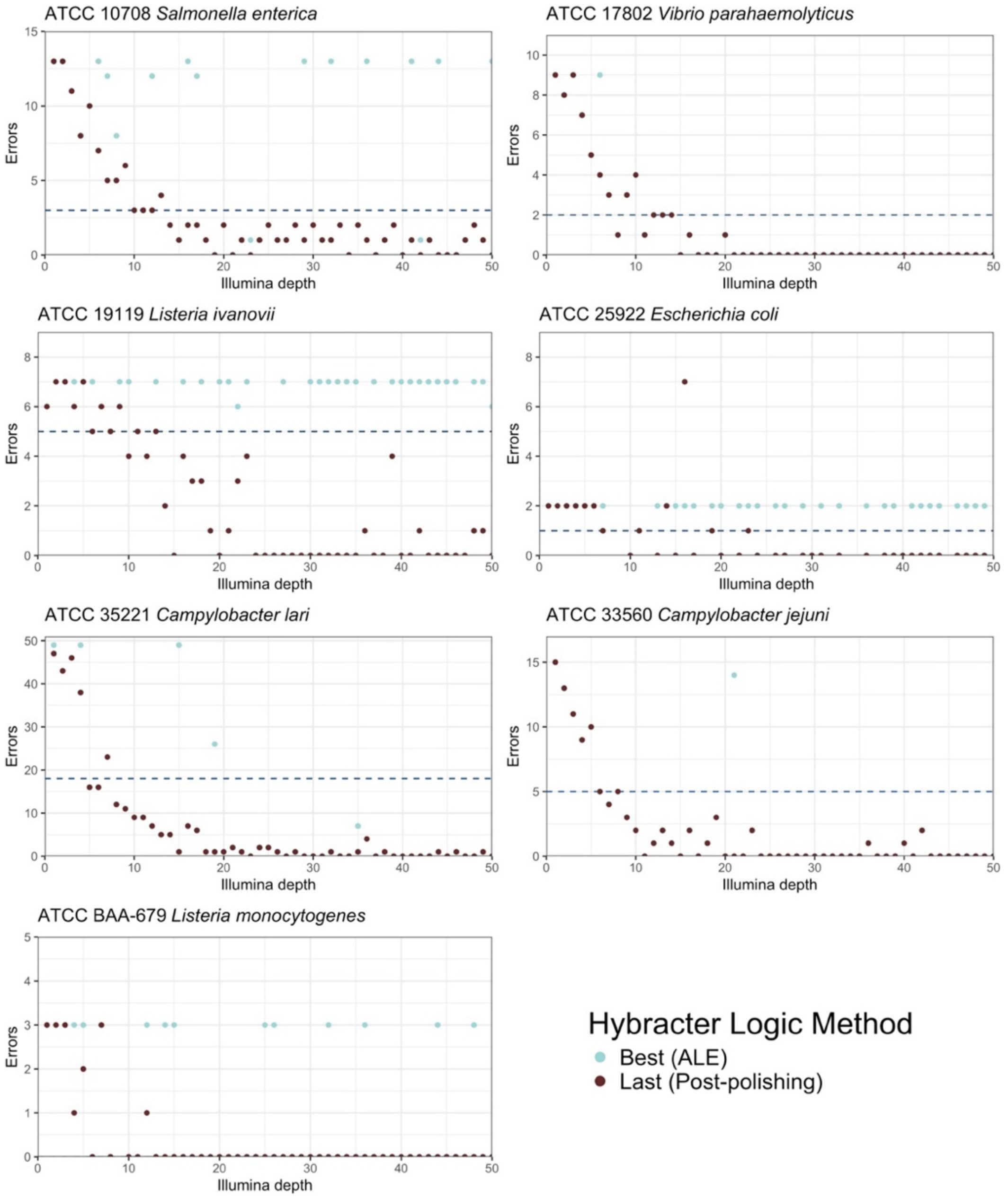


**Figure S7: Hybracter v0.7.0 with short-read polishing errors per genome chosen by ALE score.** For each of the seven reference genomes without errors and significant non-determinism in their automated long-read assemblies, this figure shows the number of errors remaining after assembly with Hybracter v0.7.0 using the parameters '--logic best' or '--logic last'. Each genome was tested at each interval from 1× to 50× depth (x-axis). The dashed blue lines represent the Trycycler long-read only assembly error count for each genome, with ATCC BAA-679 *L. monocytogenes* having 0 errors in its Trycycler long-read only assembly. All blue points with '--logic best' are where Hybracter takes the polishing round with the highest ALE score as the 'best' final assembly.  All maroon points with '--logic last' take the final short-read polishing round as the best final assembly regardless of ALE score. If best and last methods provide identical results, only the maroon last points are plotted. As can be seen, the best ALE score polishing round never outperforms taking the final polishing round for any genome other than one instance in *E. coli* (at 16×), and for some genomes (*L. monocytogenes*, *L. ivanovii*, *E. coli*, *S. enterica*) is frequently worse. From v0.7.0 Hybracter therefore implements '--logic last' by default and deprecates '--logic best'.
